# *In-silico* Structural and Molecular Docking-Based Drug Discovery Against Viral Protein (VP40) of Marburg Virus: A Causative Agent of MAVD

**DOI:** 10.1101/2021.01.26.427942

**Authors:** Sameer Quazi, Tanya Golani, Nashat Akhta, Christina Elsa Thomas, Zeshan Haider

## Abstract

The Marburg virus (MARV) is reported to induce extreme hemorrhagic fever (MHF) with a high degree of infectivity and lethality in both human and non-human primates. An appropriate vaccination for this virus’s treatment is not yet usable, and thus needs intensive attempts on multiple scales. In this study, we employed the Computer-Aided Drug Design (CADD) based approach to identify the drug-like compounds inhibiting the replication of the Viral protein (VP40) of MARV. Our database search using an online database “PubChem” retrieved ∼3000 compounds structure-based similarity. Lipinski’s rule was applied to evaluate further the drug-like compounds, followed by molecular docking-based screening, and the selection of screening ligand complex with VP40 based on S-score (lower than reference Favipiravir inhibitor) and root-mean-square-deviation (RMSD) value (probably less than 2) using AutoDock 4.2. Resultantly, ∼100 compounds were identified having strong interaction with VP40 of MARV. After evaluating their binding energy using the AutoDock 4.2 software, four compounds (CID-67534452, CID-72201087, CID-123273976, CID-153708661) were identified that showed strongest binding energy with VP40 of MARV and strong inhibition effect than the Favipiravir. Robust binding energy, useful ADMET parameters and drug-likeness suggest that these candidates “CID-67534452, CID-72201087, CID-123273976, CID-153708661” have tremendous potential to stop the replication of MARV, hence might lead to the cure of MAVD.

## Introduction

Depending on the epidemic, MARV is the causative agent of Marburg Virus Disease (MAVD) in humans, with case fatality rates varying from 23 to 90% (Kuhn et al., 2019). MARV is one of the genera Filoviridae family members, consisting of different viruses such as Marburgvirus, Ebolavirus, Cuevavirus, Striavirus, and Thamnovirus (Languon & Quaye, 2019; Shi et al., 2018). The Filoviruses comprised various viruses such as Marburgvirus or Ebolavirus that could be induced hemorrhagic, sometimes fatal diseases in humans and animals. In contrast, some other different Filoviruses such as Cievavirus, Striavirus, And Thamnovirus are not reported to cause disease Humans and animals. The MARV is a negative-sense RNA, and its genome is non-segmented. The non-segmented genome of MARV encoded the seven open reading frames (ORFs): nucleoprotein NP, virion protein (VP) 35, VP40, glycoprotein GP, VP24, viral polymerase L (Feldmann et al., 1992). The MARV genome has shaped the filamentous virion, with a depth of 80 nm (T W Geisbert & Jahrling, 1995).

There are two species know as MARV and Ravn virus (RAVV) that are very close to each other in genomic sequences and cause severe diseases in humans (Kuhn et al., 2013). The viral proteins of MARV have similar arrangements with viral proteins of Ebola virus. The MARV was the first filovirus discovered after outbreaks in Germany and Yugoslavia (now Serbia) in 1967 (Siegert et al., 1967). After the discovery of MARV, this virus has been sporadically found in different areas of Africa. However, an epidemic was reported in 1999 in the Democratic Republic of Congo. Due to the outbreak, the significant impact of MARV was on economic growth and human community due to its high fatality rate of 83% (Bausch et al., 2006). The largest recorded MARV epidemic happened in Angola in 2005, with 252 documented human infections with a fatality rate of 90% (Towner et al., 2006). The outbreaks of MARV have started to emerge since 2005, and again Uganda epidemic in 2007. The two cases of MVD in 2008 were reported in the United States and the Netherlands. The patients were returned after visiting from Uganda (Languon & Quaye, 2019).

The antiviral drug Favipiravir has previously been identified and reported as an effective anti-Marburg virus antiviral drug. It could be particularly promising for use in the event of an outbreak, where it could be administered orally quickly and safely, even after exposure (Zhu et al., 2018). Favipiravir antiviral drug was designated for screening ligands from an online PubChem database using structure similarity-based strategy to find similar structures. Computational processes and virtual screening play a significant role in drug designing. Virtual high throughput screening (vHTS) is a well-thought-out essential component of drug designing (Lyne, 2002). Novel cheminformatics techniques such as structure-based ligands screening from databases, molecular docking, and simulation can be used to block the catalytic site, i.e., VP40 of MARV. Using these computational biology tools, the current study was planned to identify compounds with strong interaction, greater binding energy and strong inhibition effect with VP40 of MARV. These candidates of drug-like compounds will have greater potential to stop the replication of MARV that would ultimately help development of MAVD cure.

## METHODOLOGY

### Sequence retrieval and primary structure analysis

The protein sequence MARV VP40 (Accession # APQ46231.1) was retrieved from an extensive online database NCBI (https://www.ncbi.nlm.nih.gov/). Besides, physicochemical properties to test the primary VP40 protein structure including the molecular weight of protein, amino acids composition of protein, atomic composition of protein, the extinction coefficient of protein, the approximate half-life of protein, instability index of protein, aliphatic index of protein and broad hydropathicity average (GRAVY) of protein was analyzed using the Expasy-Protparam online software.

### Homology Modeling and Evaluation of 3D Structure

The MODELLER v9.25, a homology-based desktop software, was used to predict the 3D structure of MARV’s VP40 protein (ul Qamar et al., 2020). The best structure was chosen based on the highest discrete optimized protein energy (DOPE) score of predicted ∼50 structures. Besides, the quality of the three-dimensional structure of VP40 was evaluated by using the online program PROCHECK (https://servicesn.mbi.ucla.edu/PROCHECK/).

### Preparation of Coordinate file

The protein structure of VP40 has been optimized by using Discovery Studio Visualizer and AutoDock4.2 desktop software. The structure was optimized by including various steps, such as removing water molecules from structure and crystallization with the ligand, adding hydrogen, adding the charges Gasteiger-Marsili and Kollman, non-polar hydrogen fusion, and spinning the delivery of the bonds. The script has been stored in pdbqt format for more process.

### Ligand Selection and Virtual Screening

The Favipiravir antiviral drug was screened for docking. Docking has been done with the software AutoDock 4.2. A high drug score was chosen for the catalytic domain VP40 MARV using the online tool DoGSite (https://proteins.plus/help/dogsite). The following catalytic sites were docked with antiviral drugs: “LEU84.PRO85, LEU86, GLY87, ILE88, SER90, ASN91, PHE113,GLN118,PHE120,ARG122,LEU135,ARG136,MET137,LEU138,GLU140,ASN142,GL N143,ALA144,PHE145,ILE146,MET149,VAL150,TRP179,ARG180,PRO181,LEU186,VAL19 5,SER196,VAL197,HIS198,PRO199, ILE204, VAL205, VAL273, TYR274, PHE275, GLN276, ALA277, PRO278, PHE281,GLN290,VAL292,LEU293,ALA294, TYR295, ALA296, ASN297, PRO298, SER301, ALA302, VAL303”. The following scores were calculated and picked in screening for the other antiviral compounds: Score and RMSD and binding energy score of MARV VP40 catalytic domain. The pre-eminent ligand was tested to identify drug-like compounds using online PubChem (https://pubchem.ncbi.nlm.nih.gov/) database, and the thumb rule was utilized to test the drug-like properties of structures. Various drug-likeness parameters such as molecular weight less than 500 Da, LogP less than 5, H-bonds donor less than 10 and H-bond acceptors less than 5, were considered (Lipinski, 2004). Finally, compounds that satisfy the above criteria were included in a novel database for molecular docking.

### Molecular docking

For additional assessment of these drug-like compounds, all selected compounds were docked with MARV VP40. Molecular docking simulations were carried out on an operating system on Windows 10 PC-based computers (x86). The applications included AutoDock 4.2, using the Python 2.7 -Cygwin C:\program language and Python 2.5 using MGL tools 1.5.4 (www.scripps.edu) (Morris et al., 2009). The 2D and 3D interaction analysis of docked active sites of molecules was visualized using Discovery Studio Visualizer v3.1 and PyMol v9.25 software (Inwood et al., 2009; Studio, 2008). As a receptor for docking, the energy-reduced structure was further used. The contact energies grid maps of various kinds of atoms are precalculated using AutoGrid 4.2. A grid box was built in each dock for VP40 MARV using a grid map of 45 × 45 × 45 points, 60 × 60 × 60 grid spacing points and 0.375 Å and 0.420 Å respectively. The grid maps are based on the respective binding location of the ligand inside the protein structure. The docking was conducted with chosen parameters using AutoDock 4.2: the number of GA runs was 150; the maximum number of evaluations (short) was 250000; the maximum number of generations was 27000; the number of GA runs was 10; and the gene mutation rate was 0.02 to define and test the relationship between ligands and VP40 (Mujwar & Pardasani, 2015). The S value (negative) is a scoring value that calculates the ligand’s receptor affinity and is determined by the AutoDock 4.2 default scoring-built feature. RMSD is used to equate docked conformation with reference-docked conformation. As a possible barrier, the recovered compounds may be produced that have lower S-score and RMSD value than the reference substrate (Haider et al., 2020). The binding energy of these hits was established for further study. The binding affinity revealed the ligand’s hydrophobic and polar relationship with the active receptor site. Its 5 to 15 kcal/mol range is known to be a potent ligand-receptor interaction. A compound with a binding affinity like or greater than that of the Favipiravir inhibitor is known for further study of ingestion, excretion of delivery metabolism, and toxicity profiling. The physical and chemical properties of drug-like compounds have been analyzed using the AdmetSAR server.

### AdmetSAR Profiling and Toxicity Validation

The level of toxicity of selected compounds was evaluated by using an online tool admetSAR (Immd.ecust.edu.cn/admetsar2). This online software predicts the different toxic effects such as mutagenicity, irritation actions, mutagenicity, productiveness. The properties of drug-likeness and goof ADMET profile help select safe antiviral drugs for humans (Monteiro et al., 2019).

## RESULTS

The numerous research studies have been carried to find the therapeutic vaccine against MARV, but there is no drug/vaccine available to efficiently combat this pathogen (Bozhanova et al., 2020). Therefore, it is necessary to find a cost-effective antiviral drug that controls MARV. An efficient technique for drug discovery is to test whether existing drug-like compounds are effective against viral infections. The conventional drugs discovery methods are time-consuming, laborious, and less efficient (Qamar et al., 2016). Considering overhead discussion, the present study mainly focused on the structure-based virtual database screening, molecular docking and drug-likeness profiling—the selected compounds with strong binding effect with VP40 catalytic site of MARV.

### Sequence retrieval and primary structure analysis

The MARV VP40 protein sequence of 303 amino acids was retrieved from NCBI database. The protein structure viability relies on the three-dimensional conformation of the target protein. Based on their physicochemical properties, protein sequences have been constructed. The physicochemical properties predicted by Expasy-program showed that the VP40 viral protein of length 303 AA with a molecular weight of 106,660.24 Da, a GRAVY score of − 0.296, an index of the instability of 27.03, and can form hydrogen bonds, classifying it as a particular protein (Table 1).

**Table 1.**
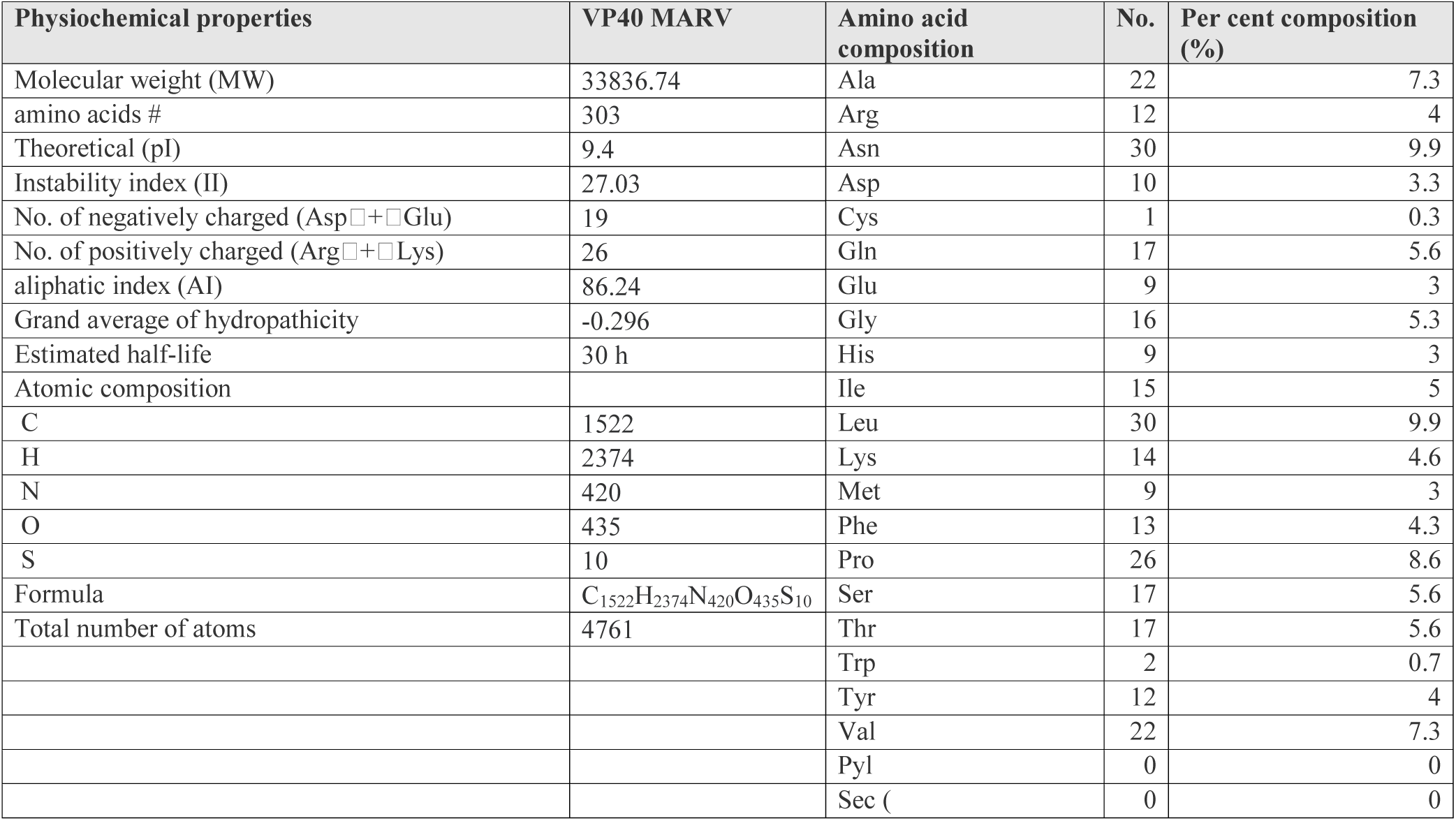
Physiochemical properties of VP40 protein

### Homology Modeling and Evaluation of 3D Structure

The homology modelling of VP40 of MARV was completed by selecting a template “C5b0v” from PDB based on highest 93% of sequence similarity. Using the “C5b0v” protein structure as a template, 50 models were generated using a desktop software MODELLER 9.25v. The best 3D structure of VP40 was selected with a DOPE score of -29742.49609 and GA-341 score (figure 1). Ramachandran plot assessment using PROCHECK software suggests that most amino acids (215 AA) with 94.3 % are within desirable regions. Besides, the outlier of AA has been seen on the plot indicates that desired AA to be attacked by our ligands located in favoured areas. There was no need to refine and structure the study for further assessment (figure 2).

**Figure 1.**
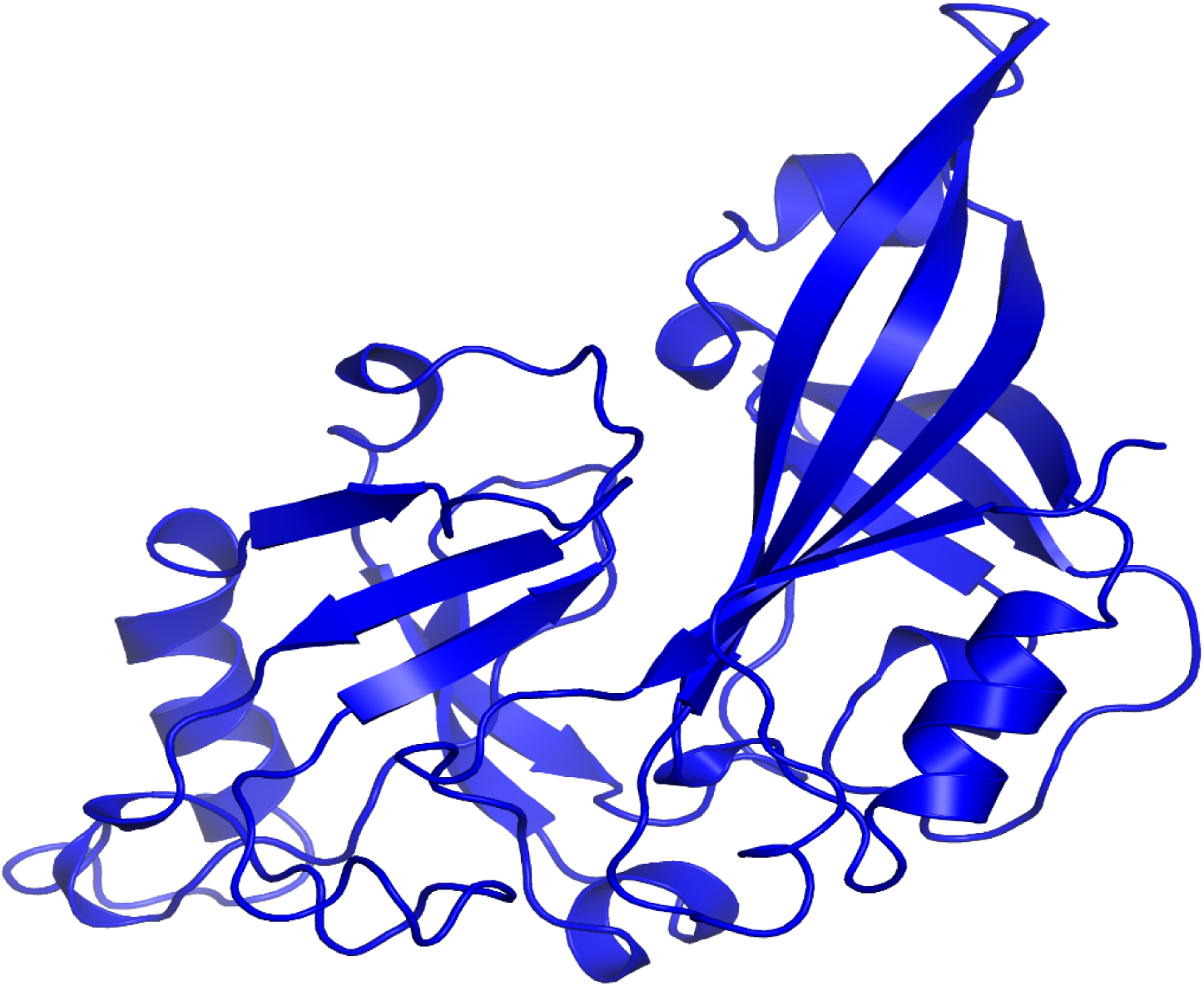
3D structure of VP40 of MARV by utilizing a template (C5b0v) and represented using PyMol v9.19.

**Figure 2.**
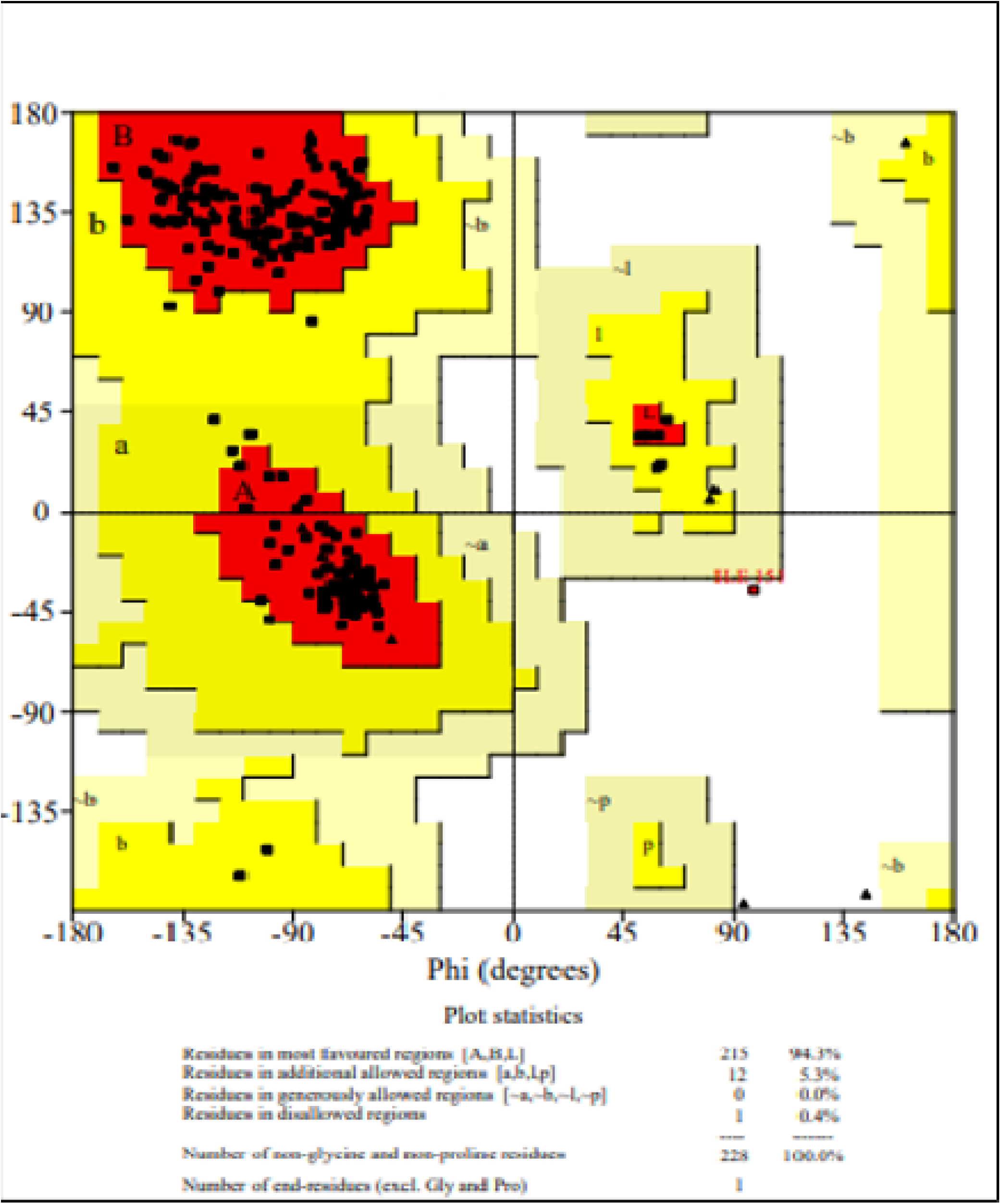
Ramachandran’s plot assessment of MARV VP40 protein represents that 94.3% residues are presented at favored regions, while ∼5.3% residues are in allowed areas, and 0.4% residues are presented outlier regions.

### Ligand Selection and Virtual Screening

The VP40 protein of MARV was docked with the selected ligand from a previous literature survey. Table 2 displays the docking score, RMSD, and binding energy of ligands with the VP40 receptor. The findings showed a good association of Favipiravir with the VP40 of MARV (figure 3). Therefore, for simulated screening, we selected the Favipiravir with S-score (− 10.9714), RMSD (1.5514) and binding energy (− 11.322). For ligand-based virtual screening from the online extensive PubChem database, compounds with>95 percent structural similarity to Favipiravir were selected by the following parameter. A total of 3000 combinations were retrieved, and Pfizer’s rule of five was extended to all the compounds. The 100 out of 300 compounds were arranged into a new database for docking with the target VP40 protein after energy minimizing.

**Table 2:**
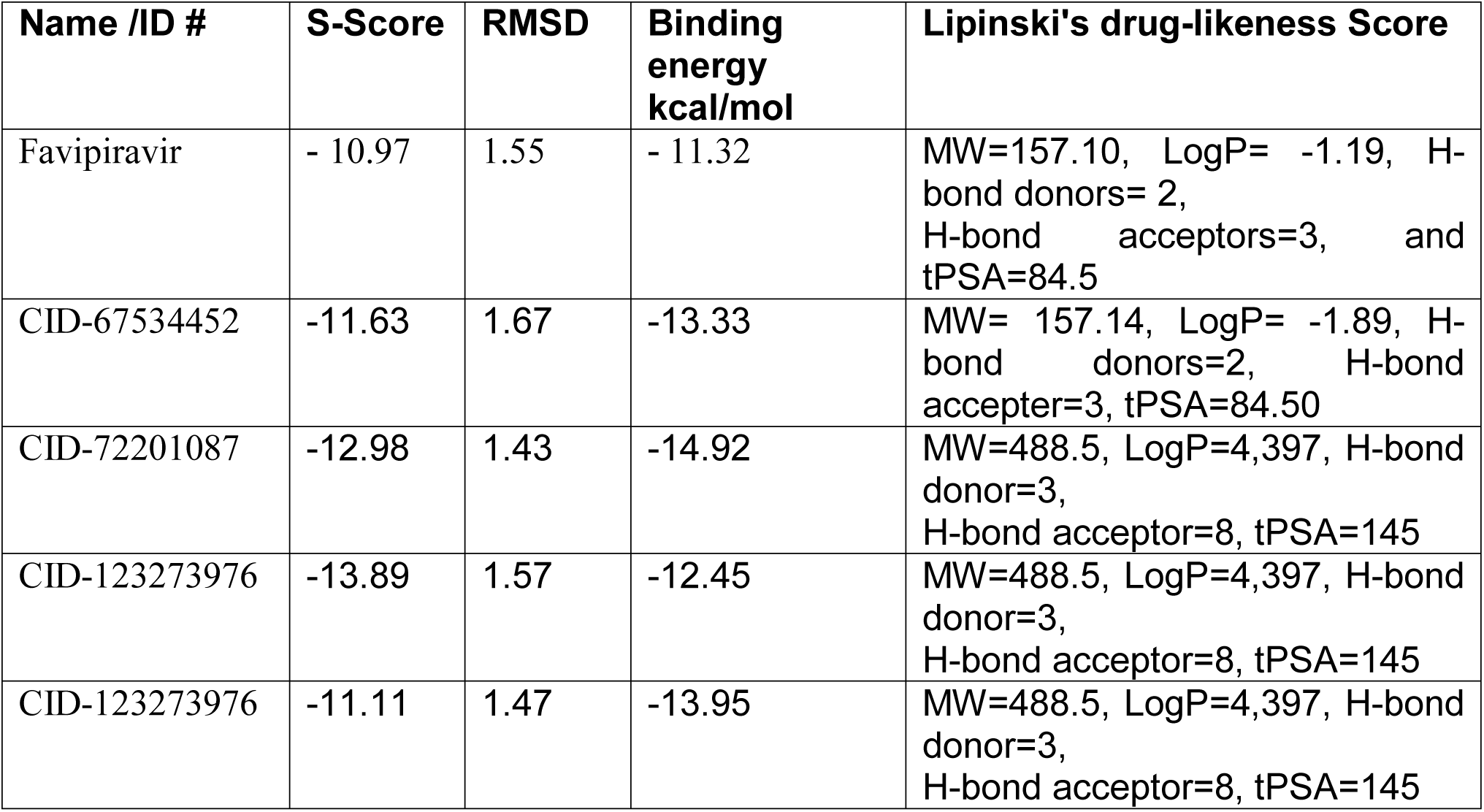
The best docking score, RMSD, and Lipinski’s rule scan results of the selected compounds.

**Figure 3.**
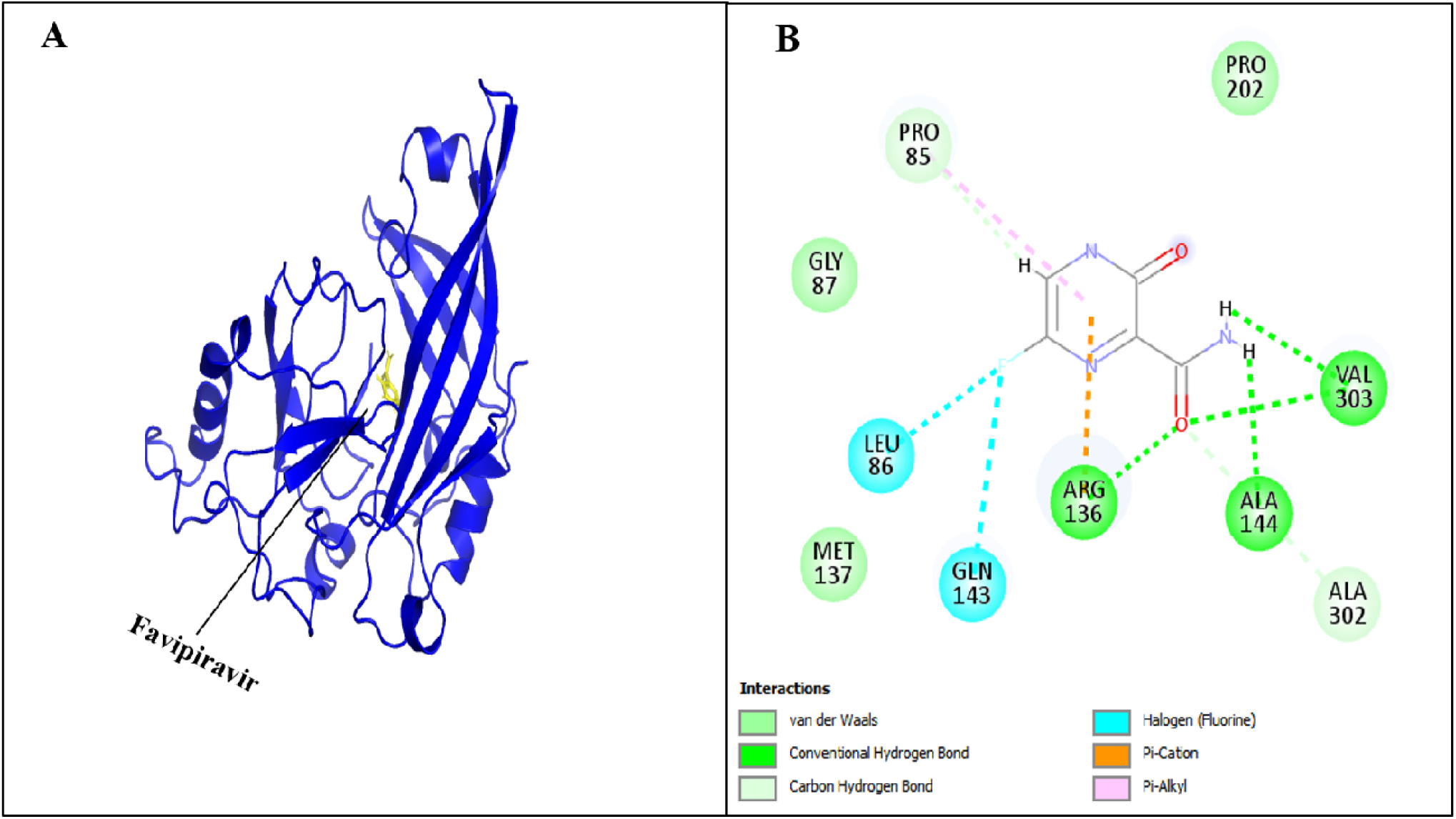
**(A)** 3D representation of VP40 MARV (Blue colour) docked posed with most active binding pocket’s residues of “Favipiravir” (Yellow colour) **(B)** 2D model of ligand “Favipiravir” compulsory mode with receptor active site of VP40 MARV.

### Molecular docking

The importance of molecular docking is commonly known for developing medicines against numerous fatal diseases. Using the Auto Dock 4.2 docking algorithm, all the hits were docked against VP40’s most successful location (P1). Consequently, four compounds with a lower S-score (docking score) than Favipiravir have been reported. For further assessment, the better docked hit compounds having lower S-score and RMSD value were chosen. Using AutoDock 4.2 tools, the binding interaction of these four hit compounds with VP40 was calculated. Based on the lowest binding energy in the largest cluster, the number of hydrogen bonds with active site residues and the protection of interactions with those from control docking, the best positions were described in the prescribed order of choosing (table 2). To guarantee that the hits docked precisely in the right binding place of importance, this was achieved. The impacts that demonstrated a significant association with MARV VP40 active site are successful inhibitors (figure. 4).

**Figure 4.**
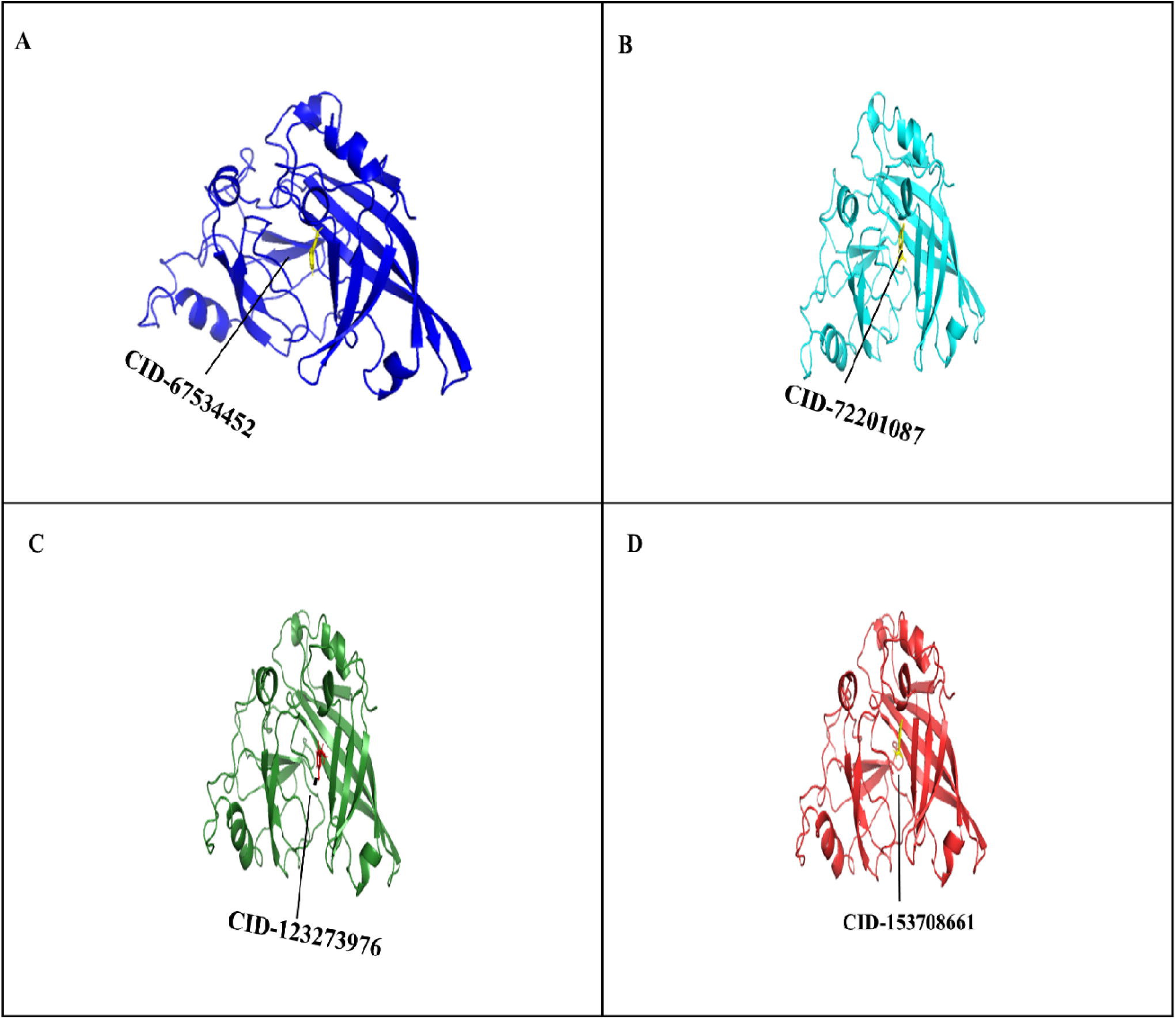
Receptor-ligands (CID-67534452, CID-72201087, CID-123273976, CID-153708661,) binding interaction analysis. **(A)** shows the 3D computationally predicted structure of MARV VP40 (Blue colour) complex with novel inhibitor CID-67534452 (Yellow colour) **(B)** shows the 3D computationally predicted structure of MARV VP40 (Cyans colour) complex with novel inhibitor CID-72201087 **(C)** shows the 3D computationally predicted structure of MARV VP40 (Green forest colour) complex with novel inhibitor CID-123273976 (Red colour). **(D)** shows the 3D computationally predicted structure of MARV VP40 (Red colour) complex with novel inhibitor CID-153708661 (Yellow colour). PyMol version 9.19 has used for these receptor-ligand complex. PyMol version 9.19 has been used for these receptor-ligand complex

### AdmetSAR Profiling and Toxicity Validation

To estimate the drug-like properties of proposed inhibitors against MARV VP40, the Molinspiration server was used. All the chosen compounds demonstrated a zero breach of the rule of five by Pfizer and acknowledged the drug-like properties, i.e., molecular weight (table 2). To further verify the drug-like properties, all the chosen compounds were subjected to the AdmetSAR server to predict drug likeness’s physical and chemical properties (Table 3).

**Table 3:**
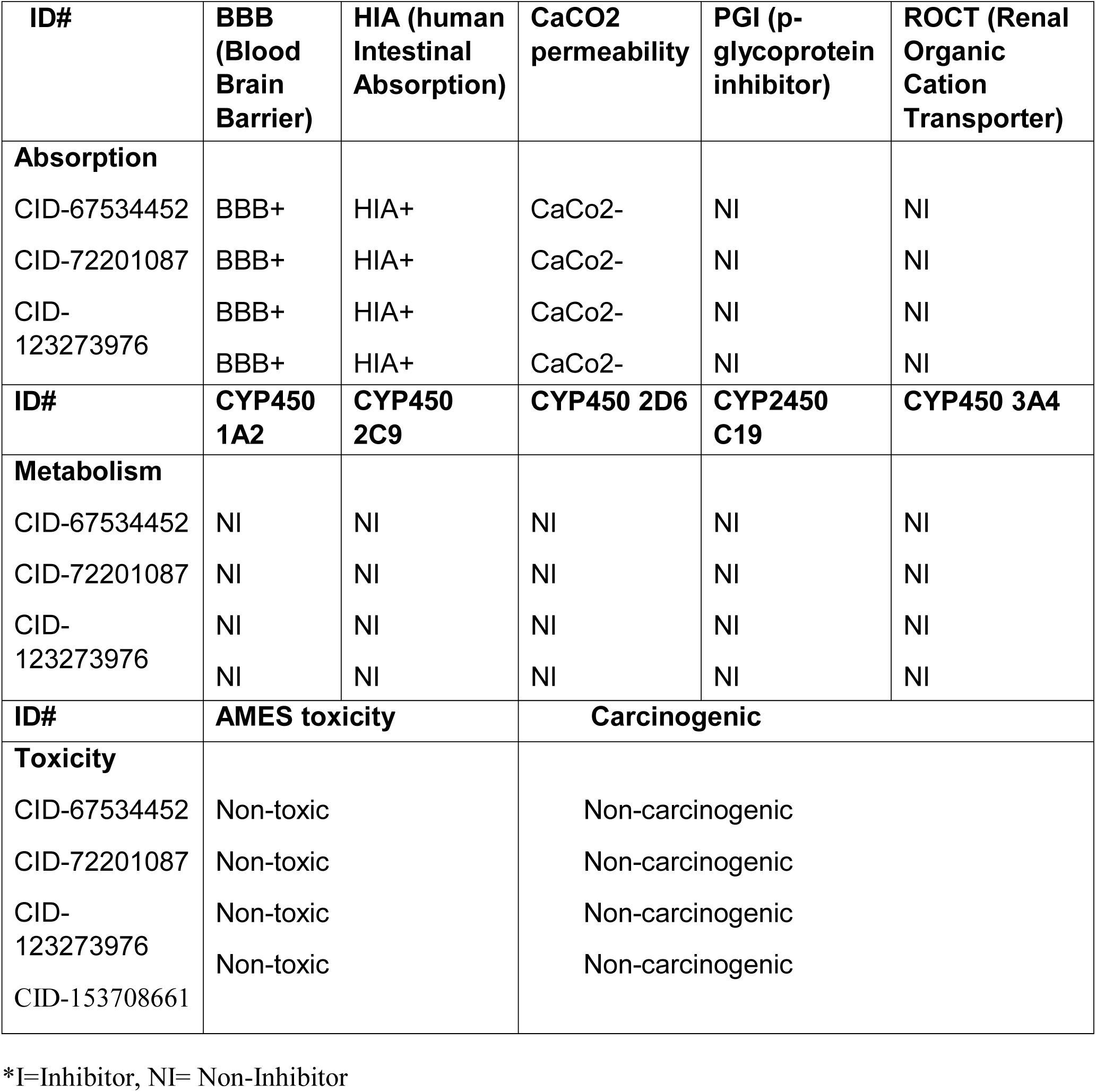
ADMET profiling for selected inhibitor’s compounds.

### Binding interaction analysis

The S-score checks the receptor’s interaction frequency with ligands. The compounds on an excellent drug compound which be selected based on the S-score, and binding energy. The following compounds CID-67534452, CID-72201087, CID-123273976, and CID-153708661, have strong interactions with the MARV’s active sites VP40. The results showed that VAL-303 and ALA-144 are most active acids from the active site of VP40, which formed the hydrogen bonding with inhibitors. While the amino acids MET-137, ARG-136, LEU-86 are amino acids that involve hydrogen bonding and carbon and polar interaction with inhibitors. ASN-142 and GLN-143 are the amino acids that took part and played a major role in forming polar interaction with novel inhibitors. The 2D and 3D docking conformations of the selected compounds are shown in Figure 5.

**Figure 5.**
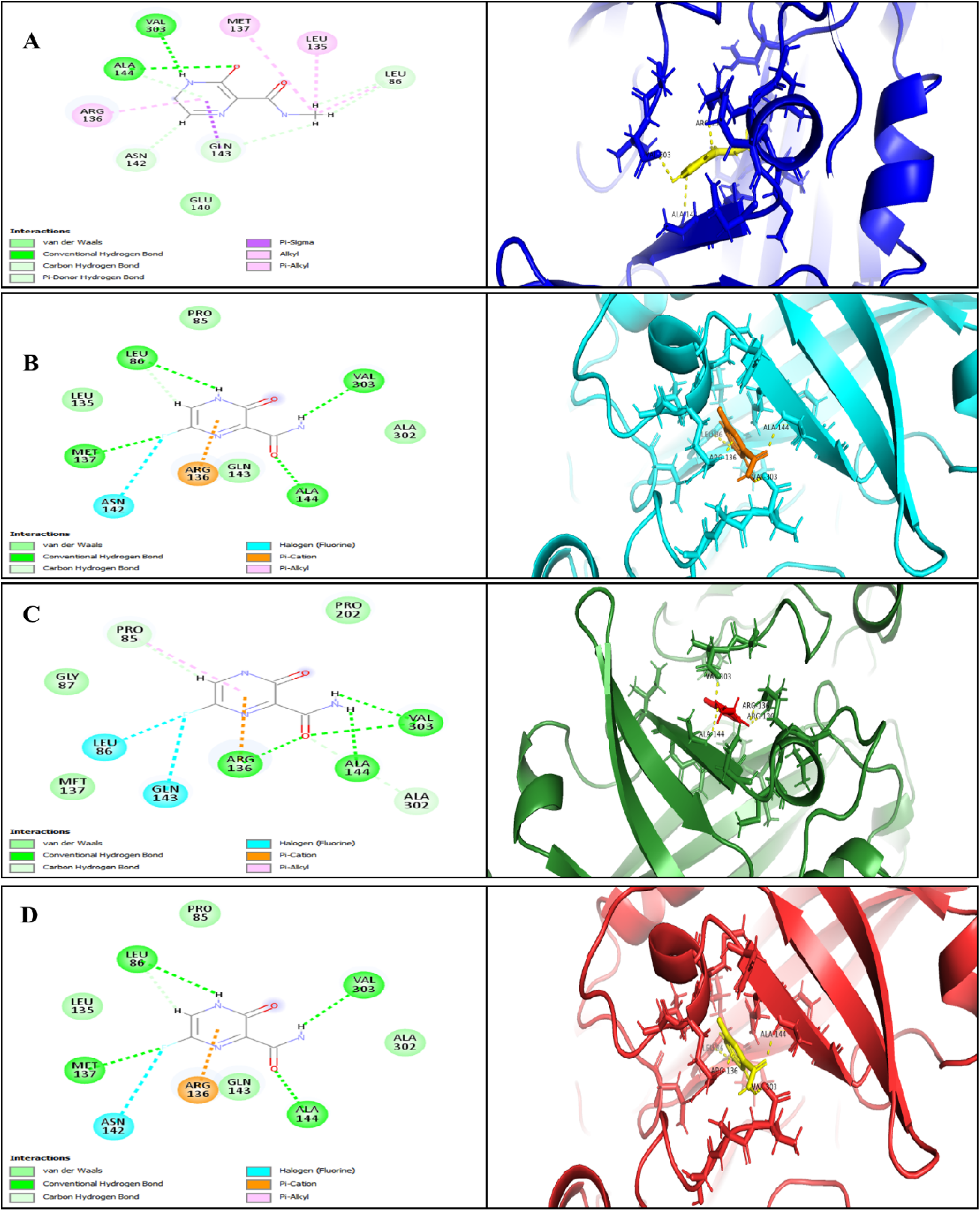
**(A)** Docked “CID-67534452” complex with VP40 protein. The amino acid residue VAL-303, ALA-144, are making hydrogen bonds. While MET-137, LEU-135, ARG-136, ASN-142, GLN-143, are forming corban and Pi-donor hydrogen bonds. **(B)** Docked “CID-72201087 complex with VP40 protein. The amino acid residues VAL-303, ALA-144, MET-137, LEU-86, are making hydrogen bonds. While ARG-136, and ASN-142, are forming corban and Pi-donor hydrogen bonds. **(C)** Docked “CID-123273976” complex with VP40 protein. The amino acid residues VAL-303, ALA-144, and ARG-136, are making hydrogen bonds. While GLN-143, LEU-86, and PRO-85, are forming corban and Pi-donor hydrogen bonds. **(D)** Docked “CID-153708661” complex with VP40 protein. The amino acid residues VAL-303, ALA-144, MET-137, and LEU-86 make hydrogen bonds. At the same time, ARG-136 is forming corban and Pi-donor hydrogen bonds.

## Discussion

Several studies have recently been carried out to find the therapeutic vaccine against MARV but with little success (Fu et al., 2016). Considering the widespread virus and the consequent human fatalities reported worldwide, there is an urgent need to find a cost-effective antiviral drug to control this menace.

An efficient technique for drug discovery is to test whether existing drug-like compounds are effective against viral infections. However, the conventional drug discovery techniques are time-taking and less efficient (Thomas W Geisbert et al., 2010). In silico study, the MARV receptor binding sites of glycoprotein are promising candidates for epitope-focused vaccine design to induce neutralizing antibody against MARV (Bozhanova et al., 2020). Considering the discussion mentioned above, the present study suggested compounds that strongly bond with the VP40 of MARV catalytic site by inhibiting its proteolytic activity and can be regarded as drug compounds. The three drug-like compounds were further evaluated with AdmetSAR server to screen ADMET-properties (Cheng et al., 2012). Blood-Brain-Barrier (BBB) is the resister present within endothelial cells and stops the brain from taking any pharmaceutical. BBB is an essential factor in the field of drug discovery (Alavijeh et al., 2005). Oral bioavailability is considered a necessary element for selecting an active drug for patients (Thomas et al., 2006). ADMET characteristics of useful drug-like compounds such as P-G substrate (P-Glycoprotein substrate), BBB penetration, HIP (Human intestinal preparation), ROCT (Renal Organic cation transporter), CaCO2 permeability, represented the positive results for the likeness of an effective drug. CYP (Cytochromep450) is a group of isoenzymes involved in the catabolism of various chemicals such as steroids, drugs, carcinogens, bile acids, etc. A profile of ADMET test for a useful and effective drug compound consisted following parameters: (a) it must pass the BBB, (b) absorb in the human intestine, (c) absorb the CaCO2 permeability, (d) non-toxic, (e) non-carcinogenic, and (f) non-inhibitor to CYP enzyme. The four (CID-67534452, CID-72201087, CID-123273976, and CID-153708661) compounds significantly accepted these ADMET parameters (table 2). The selected compounds for drug, i.e., PubChem ID, docking score, RMSD value, binding energy value, the binding residue of the VP40 of MARV, Lipinski’s results and ADMET analysis. The selected compounds are capable to serve as the novel, structurally different and potentially active inhibitors against the VP40 of MARV.

Our in-silico study found three inhibitors with strong inhibition potential effect of drug lead may be therapeutic inhibitors against VP40 of MARV by efficiently targeting the active site of VP40 of MARV. Therefore, our finding of three druglike compounds, i.e., “CID-67534452”, “CID-72201087”, “CID-123273976”, and “CID-153708661” require further in-vitro and in-vivo work for structure-based optimization before commercialization.

## Conclusion

The current study’s primary aim was the structure-based virtual screening using the PubChem database, the molecular docking of selected compounds, and the evaluation of binding interaction against the VP40 of MARV. The compounds from the online PubChem database (CID-67534452, CID-72201087, CID-123273976, and CID-153708661) showed strong interaction against the active site of the MARV VP40. Results show that these compounds could be hypothetically used as a drug against MARV. To validate these results, in-vitro and in-vivo analyses must require turning these possible inhibitors into therapeutic medicines. We anticipate that the current study’s insights may prove useful for further exploration and production of novel natural therapeutic agents against MAVD.

